# Interpreting GC content differences across populations at polymorphic sites

**DOI:** 10.64898/2026.05.16.725686

**Authors:** Sheel Chandra, Ziyue Gao

**Author notes:** Author for Correspondence*: Ziyue Gao, Department of Genetics, University of Pennsylvania Perelman School of Medicine, Philadelphia, PA, USA.

## Abstract

Recent studies have reported consistent inter-population differences in GC content at polymorphic sites in multiple species, including humans. Specifically, populations that experienced recent bottlenecks exhibit lower average GC content at common polymorphic sites compared to non-bottlenecked groups—an observation previously interpreted as an indication of rapid evolution of base composition or shifts in mutation patterns. In this study, we investigate the evolutionary and technical factors driving these patterns across humans, mice, maize, and silkworm. In humans, the reported inter-population difference at common variants reverses among rare variants and becomes nearly negligible when all polymorphic sites are considered. Including rare variants similarly reduces the inter-population differences in other species examined. Across species, we observe that rare variants consistently exhibit higher GC content than common variants, reflecting differences in the relative abundance of mutation types across allele frequencies. We demonstrate that these observations arise from the combined effects of demographic history and an excess of GC→AT over AT→GC mutations, which is counteracted by GC-biased gene conversion (gBGC) over long evolutionary timescales. Forward-in-time simulations incorporating these processes and realistic parameters recapitulate the observed variation in GC content across populations and allele frequency bins. Overall, our findings show that the GC content at polymorphic sites is strongly shaped by the interaction among demographic history, GC→AT versus AT→GC mutation imbalance, and gBGC, rather than reflecting stable, genome-wide trends. Consequently, inter-population differences in GC content—especially at common variants—should not be interpreted as evidence of ongoing divergence in base composition or population-specific shifts in mutation patterns.

**Significance Statement:** Inter-population differences in GC content at polymorphic sites in humans and several other species have been interpreted as evidence of rapid evolution in base composition— a surprising claim given the conservation of genomic GC content across species. Here, we demonstrate that these allele frequency-dependent patterns can arise predictably from the interplay between AT-biased mutational input, GC-biased gene conversion, and demographic history, without requiring population-specific changes in mutation patterns. This work provides a mechanistic explanation for a puzzling pattern observed across diverse taxa and cautions that allele frequency filters can systematically distort the inference about mutation patterns from segregating variants.

## Introduction

DNA base composition, a fundamental attribute of the genome, can vary across species but can also be surprisingly conserved between some distantly related species (Bernardi, 2000; Šmarda et al., 2014; Long et al., 2018). Understanding the evolutionary basis of these patterns can help shed light on several related genomic features, such as codon usage, methylation, and genome organization (Costantini et al., 2009; Tillo & Hughes, 2009; Plotkin & Kudla, 2011). The evolutionary forces governing these patterns have been a subject of heated debate, centered on whether genomic GC content (GC%) variation is primarily shaped by neutral mutational processes or by non-neutral forces such as natural selection.

Conceptually, the GC content of a genome is shaped by a balance of conversions between strong nucleotides (C or G) and weak nucleotides (A or T) across generations. Specifically, this is determined by both the mutation rates (⍰) and the fixation probabilities (⍰) of GC→AT and AT→GC mutations. For a genome at GC% equilibrium, the number of GC→AT substitutions should balance the number of AT→GC substitutions:

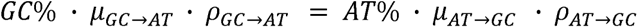

When the left-hand side of the equation is greater, more GC→AT substitutions will be fixed into the genome, reducing the overall GC content and hence the value of the left-hand side. Conversely, when the right-hand side is greater, the genomic GC content will increase until the balance is achieved.

Single nucleotide mutation rates strongly depend on the mutation type and sequence context (Hwang & Green, 2004; Lynch, 2010). Typically, the average rate of GC→AT mutations is higher than that of AT→GC mutations, referred to as mutation bias hereafter (Lynch, 2010). This GC→AT mutation bias can be partially attributed to the hypermutability of methylated cytosines but is also observed in species lacking DNA methylation (Petrov & Hartl, 1999).

Intriguingly, even in terms of mutation counts, many species exhibit a consistent excess of GC→AT mutations over AT→GC mutations, which has been observed in natural polymorphisms, *de novo* mutations, or mutation accumulation experiments (Keightley et al., 2009; Hershberg & Petrov, 2010; Bergeron et al., 2023). Hereafter, we refer to this excess in mutation counts as “mutation imbalance”. This universal imbalance suggests that mutational input alone would drive genomes toward a more AT-rich state, implying either that these genomes are far from GC equilibrium or that a compensatory fixation bias (⍰_AT_→GC > ⍰_GC_→AT) counteracts the mutational pressure.

Indeed, the probability of fixation for GC→AT and AT→GC alleles can differ due to differences in the direction, strength, and rate of natural selection as well as biased gene conversion. In genomes with large fractions of non-coding regions, differences in average fitness effects between GC→AT and AT→GC variants are likely negligible. However, GC-biased gene conversion (gBGC) is widespread across diverse taxa, including yeasts, plants, fish, birds and mammals, significantly influencing genetic variation patterns and GC content evolution (Mancera et al., 2008; Duret & Galtier, 2009; Muyle et al., 2011; Nabholz et al., 2011; Figuet et al., 2015; Glémin et al., 2015; Pouyet et al., 2018; Boman et al., 2021). gBGC occurs due to the biased repair of mismatches in heteroduplex recombination intermediates during meiosis, leading to a slight over-transmission of GC alleles over AT alleles in one generation and a greater fixation probability for AT→GC variants over evolutionary timescales. Notably, the effects of gBGC on the fixation probabilities of different mutation types depend on recombination rate, transmission bias, and effective population size (Duret & Galtier, 2009).

Recently, a study (Li et al., 2015) reported striking differences in GC% at polymorphic sites between populations within the same species. Specifically, individuals in groups that experienced recent bottlenecks and/or domestication (referred to as the “derived groups” in the original study) showed lower GC% at single nucleotide polymorphisms (SNPs) with minor allele frequency (MAF) above 5% compared to those in non-bottlenecked groups (referred to as the “ancestral groups” in the original study). This pattern is consistently observed across a wide range of species, including humans, mice, insects, and a variety of plants (Wang et al., 2019; Zhao et al., 2020; Gou et al., 2024), which was interpreted as an indication of rapid shifts in GC%, potentially foreshadowing divergence in base composition between populations.

To explain these observed inter-population differences, Li et al. hypothesized that bottlenecked populations may have accumulated mutations in DNA repair genes, resulting in a stronger GC→AT mutation bias. However, this explanation seems unlikely, at least for humans, for several reasons. First, known mutation-spectrum differences among populations are modest and context-specific, and they are unlikely to substantially alter the overall GC→AT mutation bias (Harris, 2015; Harris & Pritchard, 2017; Mathieson & Reich, 2017; Gao et al., 2023). Second, the strongest signal of mutation spectrum shifts, a European-specific TCC>TTC mutation pulse, is not found in East Asians or Americans and does not match the African versus non-African differences in SNP GC%. Finally, new mutations that arose after population split are expected to be rare and population-private, and unlikely to contribute substantially to globally common SNPs (Biddanda et al., 2020). Another possibility openly acknowledged by Li et al. is differential fixation of variants between bottlenecked and non-bottlenecked populations due to differences in demography or other evolutionary forces, possibly in interaction with the AT-mutation bias; however, the specific evolutionary mechanisms (i.e., drift, gene conversion, or selection) were not further investigated, nor was how population demographic history could influence (or correlate with) these evolutionary forces.

In this study, we investigated whether the observed inter-population differences in GC content are robust and sought to identify the technical factors or evolutionary forces driving these patterns. Our analysis reveals that GC% at polymorphic sites varies strongly with allele frequency, and consequently, inter-population GC% differences are highly sensitive to the allele frequency threshold applied—a pattern we consistently observe across human, mouse, maize, and silkworm. When the allele frequency cutoff is relaxed, the inter-population differences in GC content are drastically reduced across all species studied, reflecting opposing trends between rare and common variants. We propose a parsimonious model in which the observed differences emerge from the interaction between a universal excess of GC→AT mutations, GC-biased gene conversion, and population-specific demographic impacts on the site frequency spectrum. Overall, our findings demonstrate that common variants are strongly shaped by demographic history and are not representative of genome-wide patterns in base composition.

## Results

### GC content differences between human populations depend strongly on allele frequency threshold

We first replicated the observed differences in GC% among human populations using the high-coverage whole genome sequencing in the updated 1000 Genomes Project dataset (Byrska-Bishop et al., 2022). To minimize the effects of direct purifying selection, we restricted our analysis to putatively neutral regions by excluding conserved and coding sequences. To avoid population classification ambiguity, we excluded individuals from populations with clear admixed continental ancestries (population labels ACB, ASW, PUR, CLM, and PEL; see Methods). After filtering, the dataset contained 2064 individuals (504 AFR, 503 EUR, 504 EAS, 489 SAS, 64 AMR). We then calculated the global and population-specific minor allele frequency (MAF) for each SNP.

We followed Li et al.’s approach and quantified each individual’s base composition at qualifying SNPs in three steps. First, we defined qualifying SNPs based on the global MAF (specific thresholds varied depending on the analysis) in the entire dataset of 2064 non-admixed, unrelated individuals (see Methods). Next, for each individual, we tallied the occurrences of A, C, G, and T alleles on the reference strand at these qualifying SNPs based on their genotype values and allele states. For example, a homozygous T/T site in an individual contributes two counts toward the aggregate T count, while a heterozygous C/A site contributes one count to C and one count to A. Finally, we determined each individual’s overall base composition at qualifying SNPs by summing the total counts of A, C, G, and T, and plotted the fraction of C alleles against the fraction of A alleles. Because this procedure relies solely on observed allele counts of each individual, the resulting composition estimates are mathematically invariant to the choice of reference genome or ancestral polarization.

Following Li et al., we first examined common variants with global MAF above 5% (2,673,086 qualifying SNPs total). Our analysis replicated the finding of higher GC% in African individuals at common SNPs compared to non-African individuals (Figure 1A, 52.89% vs. 52.27-52.32%; Wilcoxon rank-sum test, p-value < 10^-80^). In their original study, Li et al. tested the robustness of their observation by applying different cumulative MAF filtering (≥5%, ≥1%, and ≥0.5%). They observed that inter-population differences weakened at lower MAF thresholds and attributed this attenuation to potential contamination from sequencing errors or somatic mutations (Li et al., 2015, Supplementary Figure 5). Consistent with this prior observation, when we recalculated GC content across progressively lower MAF cutoffs, the inter-population gaps gradually shrink, decreasing from ∼1% at MAF≥10% to ∼0.008% without MAF filtering (Figure 1B, 55.474% vs 55.466% for African and non-AFR, respectively; Wilcoxon rank-sum test, p-value < 10^-80^), leaving only a minute absolute difference despite high statistical significance driven by the large number of variants.

**Fig. 1.**
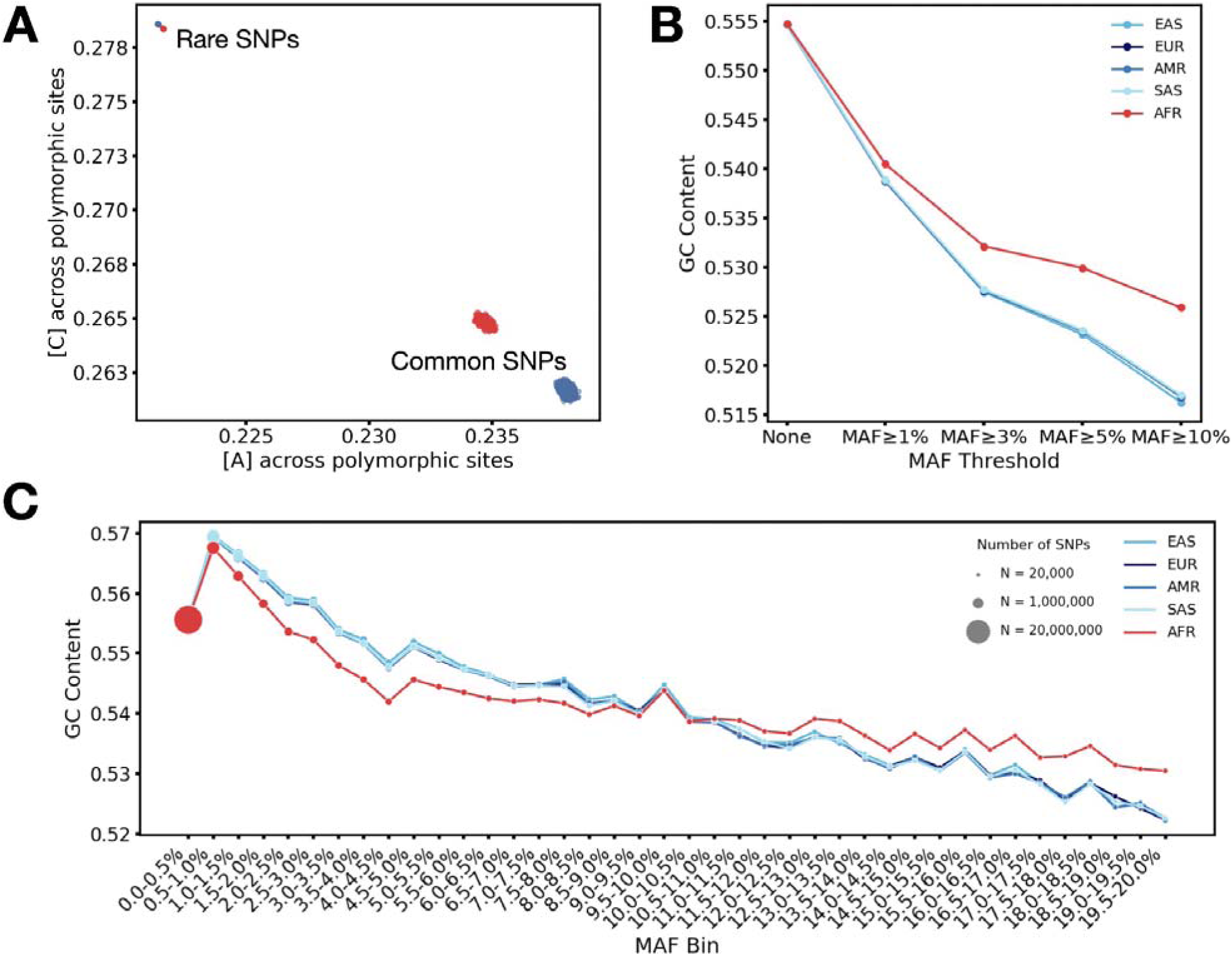
GC content across polymorphic sites is dependent on MAF filter applied. **A.** Reference strand base composition plotted as proportion of A vs. proportion of C across common polymorphic sites (MAF≥5%) and rare polymorphic sites (MAF<5%) in AFR (red) and non-AFR (blue) groups. Individuals in populations with admixed ancestries (i.e., with population labels ‘ACB’, ‘ASW’, ‘PUR’, ‘CLM’, ‘PEL’) are excluded. **B.** GC% across polymorphic sites calculated from different variant sets: No MAF threshold, ≥1%, ≥3%, ≥5%, and ≥10% MAF for each of the 5 population groups in 1000 Genomes. Each population group is represented by a unique color, with AFR in red and non-AFR groups in different shades of blue. **C.** population-specific GC% across SNPs in non-overlapping MAF bins ranging from 0 to 20% (0.5% width). Marker size is proportional to the number of SNPs in each bin. **Alt-text:** Plots showing GC content at polymorphic sites in different bins based on minor allele frequency (MAF), with subfigures labelled from A to C.

Critically, however, this cumulative MAF filtering did not separate rare variants (MAF<5%) from common variants (MAF≥5%). When isolated, rare variants (MAF<5%) displayed substantially higher overall GC content and a reversed inter-population pattern: African individuals have lower GC content than non-African individuals (Figure 1A, 55.65% vs. 55.69%; Wilcoxon rank-sum test, p-value < 10^-80^). This reversal offers an alternative explanation for the attenuated gap observed at lower MAF cutoffs: rather than technical noise diluting a genuine signal, opposing biological effects of rare and common variants cancel each other out in the aggregate base composition.

To distinguish between these two hypotheses and determine where this reversal occurs across the frequency spectrum, we computed GC content within non-overlapping, fine-resolution MAF bins (Figure 1C). This analysis reveals a striking crossover point at ∼10% MAF: above 10%, non-African populations exhibit a lower GC% than Africans, consistent with the trend reported for common variants with MAF≥5%; below 10%, however, the direction flips. Although we observe a sudden, aberrant drop in GC% exclusively in the rarest bin (0<MAF≤0.5%)— consistent with effects of technical artifacts—the trajectory across all remaining bins forms a continuous downward trend for each population. This continuity indicates that the reversal at around 10% MAF is a genuine biological signal related to demographic differences. Together, these results demonstrate that common SNPs represent a biased subset of segregating sites, whose apparent population divergence in GC% is largely counterbalanced by opposing trends at rare variants, markedly diminishing the overall magnitude of GC differences across all polymorphic sites.

### Including rare variants greatly reduces inter-population differences in GC content at polymorphic sites across species

To determine whether the frequency-dependence of inter-population GC% differences is a general phenomenon, we extended our analysis to three additional species: mouse (between subspecies *Mus musculus castaneus and M.m. domesticus, M.m. musculus*), maize (between subspecies *Zea mays mays* and *Z.m. parviglumis*), and silkworm (between *Bombyx mori* and *B. mandarina*). These species were previously reported to exhibit reduced GC content in bottlenecked or domesticated populations at common SNPs (Li et al., 2015; Wang et al., 2019; Zhao et al., 2020; Gou et al., 2024). Across all species examined, the magnitude of the inter-population difference depended strongly on the applied allele frequency threshold (Figure 2, Tables S1,3,4).

**Fig. 2.**
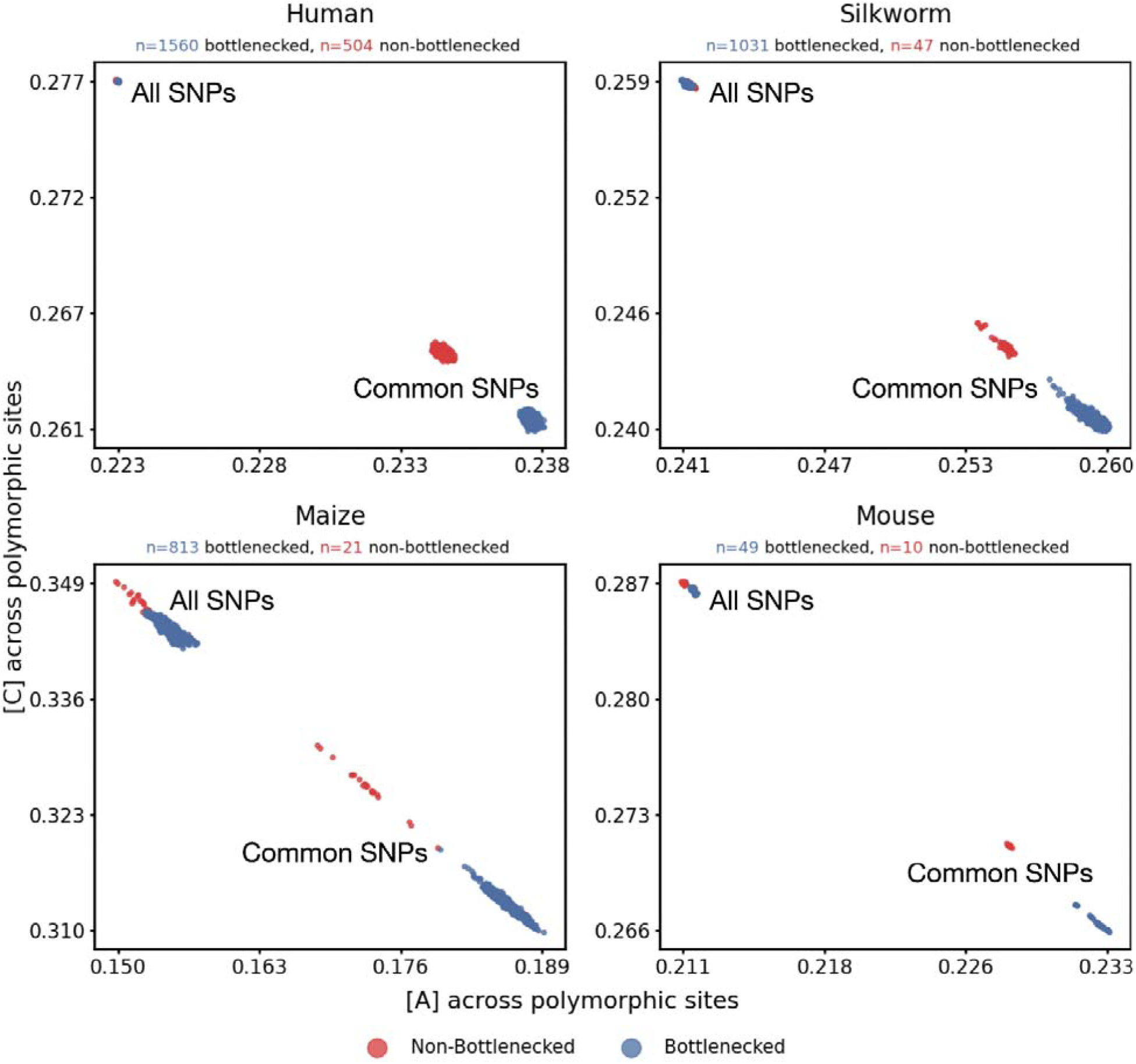
Reduced differences in GC content at polymorphic sites between bottlenecked and non-bottlenecked groups when MAF threshold is removed. Reference strand base composition plotted as proportion of A vs. proportion of C across common polymorphic sites (MAF≥5%) and all polymorphic sites (no MAF threshold) between bottlenecked (blue) and non-bottlenecked (red) populations in human, mouse, maize, and silkworm. Numbers of individuals sampled in each species are indicated on top of each panel. **Alt-text**: Scatter plots comparing GC content at polymorphic sites between non-bottlenecked and bottlenecked populations.

In all four species examined, rare variants consistently exhibited higher GC content than common variants, and crucially, inclusion of rare variants substantially attenuated the previously reported differences between bottlenecked and non-bottlenecked groups (Tables S1,3,4, Figure 2). This pattern remained qualitatively unchanged when singleton variants were excluded (Figure S1). In human and silkworm, the differences diminished to nearly negligible levels; in mouse and maize, the initial GC% differences observed at MAF≥5% were reduced in magnitude by ∼90% and 76% respectively when all variants were considered. The residual difference in mouse and maize may reflect the small sample sizes of their non-bottlenecked cohorts (n=10 and 21, respectively), which severely limit the detection of rare variants, although the non-bottlenecked group in silkworm was also relatively small (n=47). Additional sequencing across larger cohorts will be needed to confirm whether these residual inter-population differences are biological or technical artifacts. Nevertheless, the substantial attenuation of inter-population differences after inclusion of rare variants was consistently observed across all species analyzed, indicating that the sensitivity of GC content estimates to allele frequency conditioning is widespread across phylogenetically diverse taxa.

### Demographic history, GC→AT mutation imbalance, and biased gene conversion jointly shape GC content at SNPs across frequency bins

The recurrence of the threshold effect across four phylogenetically diverse species suggests that the observed GC content differences are unlikely to be driven primarily by lineage-specific changes in mutation bias. We therefore examined whether a more general mechanism— interaction between demographic history, a general excess of GC→AT mutations, and GC-biased gene conversion (gBGC)—could explain the frequency-dependent patterns. We began with human populations, where demographic history is relatively well characterized and genomic data are most comprehensive.

To generate the unfolded SFS for human populations, we determined the ancestral and derived alleles of each SNP using the inferred ancestral human genome (see Methods) and classified SNPs into six types based on the inferred mutational event that gave rise to the polymorphism: C>T, C>A, C>G, T>C, T>G, and T>A. For analyses corresponding to Figures 1-2, only minor allele frequency (MAF) was used, which is independent of ancestral-state inference, whereas analyses involving mutation type classification and derived allele frequency (DAF) require ancestral state polarization (e.g. Table 1, Figure 3). As expected under known demographic models, African populations exhibited an enrichment of intermediate-frequency variants (5%<DAF<45%) and a depletion of low- and high-frequency variants relative to non-African populations across all mutation types (Figure 3A). To minimize potential artifacts arising from recurrent mutations or mis-inference of ancestral alleles, we repeated this analysis after excluding CpG/TpG/CpA sites and obtained qualitatively similar results (Figure S2). This indicates that the observed inter-population differences in the SFS likely reflect demographic history rather than being primarily driven by recurrent mutations or ancestral allele mis-polarization.

**Fig. 3.**
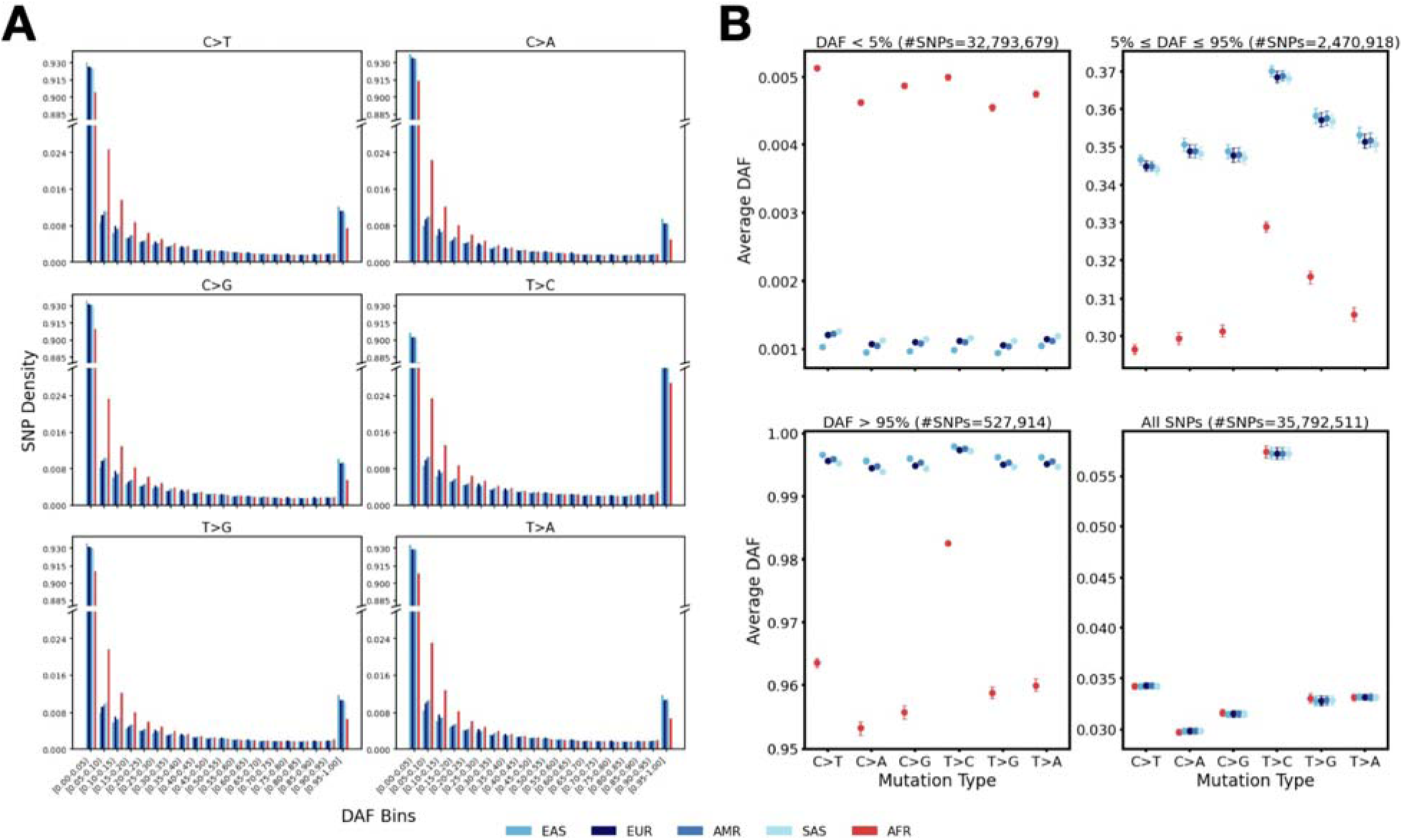
Site frequency spectrum (SFS) and average derived allele frequencies (DAF) for SNPs generated by different mutation types. **A.** SFS for SNPs of each mutation type across five human population groups. The y-axis is broken between 0.024 and 0.885 to facilitate visualization of differences at both rare and common variants. **B.** Average DAFs for each population group across different variant sets stratified by DAF: DAF<5%, 5%≤DAF≤95%, DAF>95%, and all SNPs, for each mutation type. Error bars represent 95% confidence intervals estimated using block bootstrap (see Methods). Each population group is represented by a unique color, with AFR in red and non-AFR groups in different shades of blue. **Alt text:** Two panels showing SNP density across different derived allele frequency (DAF) bins and the average DAF for each mutation type for a given population group.

**Table 1.**
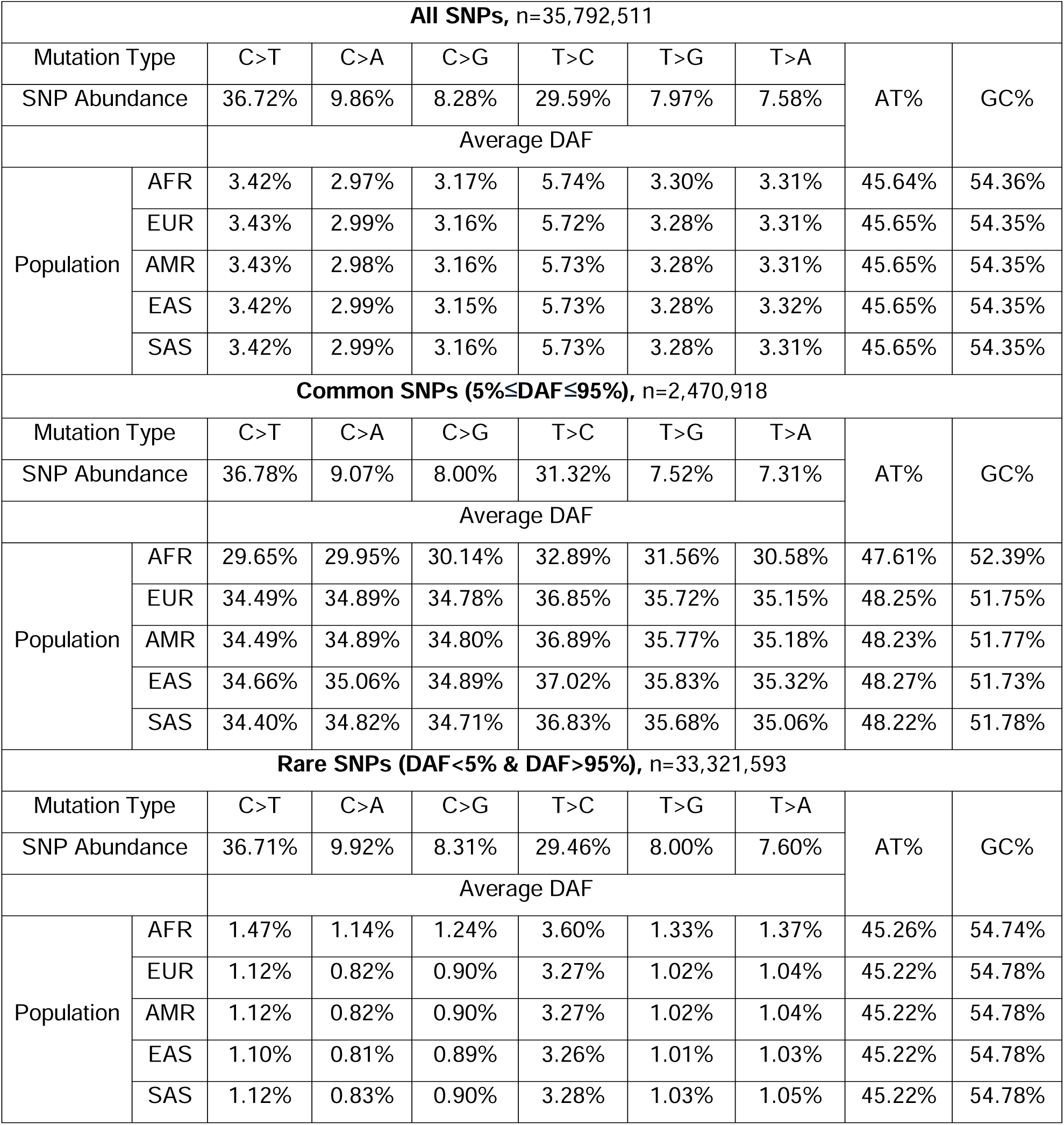
Average DAF for each population group in 1000 Genomes at a given mutation type along with total AT and GC% across SNPs in different variant bins stratified by DAF: all, common (5%≤DAF≤95%), rare (DAF<5% & DAF>95%). CpG/TpG/CpA sites are included unless indicated otherwise.

Because populations differ in their SFS, African and non-African populations also differ systematically in average derived allele frequency (DAF) in different SNP sets (Figure 3B, Table 1). For all mutation types, African populations exhibit lower average DAF than do non-AFR populations at common SNPs (i.e., 5%≤DAF≤95%) as well as at high-DAF rare SNPs (DAF>95%), whereas the opposite is true for low-DAF rare SNPs (DAF<5%) (Table S1). At the same time, GC→AT mutations outnumbered AT→GC mutations among common and low-DAF rare variants (Table S1), with qualitatively similar patterns when excluding CpG/TpG/CpA sites (Table S2). Together, these observations explain the contrasting GC content patterns across frequency bins: because derived alleles at GC→AT mutations are AT-increasing, bottlenecked populations with higher average DAFs at common variants exhibit lower GC content at those sites (Table 1).

Notably, the average DAFs across all SNPs in aggregation are remarkably similar among populations, despite substantial differences within individual frequency bins (Figure 3B). This cancellation of average DAF differences holds true across all mutation types. Consequently, the aggregate GC content across all variants differs only minimally across populations, consistent with our earlier observations (Figure 1). We also observed similar SFS shifts in mouse and maize, where bottlenecked groups displayed elevated average DAFs at common SNPs (Tables S3 and S4). These results indicate that the previously reported GC content differences at common SNPs primarily arise from demographic distortions of the SFS combined with the universal excess of GC→AT relative to AT→GC mutations observed across diverse species (Hershberg & Petrov, 2010; Lynch, 2010; Ossowski et al., 2010), without invoking population-specific changes in mutation bias.

The same framework also explains why rare variants (operationally defined as MAF<5% but dominated by variants with DAF<5%) consistently display higher GC content than common variants (MAF≥5%) across all species analyzed (Figure 1A, Figure 2, Tables S1-4) as well as why GC% decreases with MAF across the full frequency spectrum (Figure 1C). Because low-DAF rare variants by definition have a lower average DAF, and GC→AT mutations substantially outnumber AT→GC mutations in this set (Tables S1-S4), the aggregate base composition at these rare SNPs is dominated by ancestral G/C bases (the contribution of high-DAF variants to all rare variants with MAF<5% is negligible). In contrast, common SNPs have a higher average MAF, which increases the contribution of derived A/T alleles, lowering the GC% at common variants relative to rare variants.

Why, then, are GC→AT mutations more abundant than AT→GC mutations among segregating variants? A natural explanation is gBGC, which leads to fixation bias favoring AT→GC over GC→AT alleles. If the genome is already at or very close to equilibrium GC content, GC→AT and AT→GC substitutions (i.e., mutations that eventually fix) must be approximately balanced over evolutionary timescales, which requires greater new mutational input of GC→AT than AT→GC variants in the presence of unequal fixation probabilities due to gBGC.

If gBGC contributes to the inter-population differences we observe, its effect should increase with local recombination rate (Duret & Arndt, 2008; Katzman et al., 2011; Meunier & Duret, 2004; Pouyet et al., 2018). We thus stratified genomic regions by local recombination rate into quartiles (see Methods) and recalculated GC content at polymorphic sites for each population (as in Figure 1A) in the highest and lowest recombination rate quartiles. The inter-population difference in GC% at common SNPs was indeed greater in high recombination regions than in low recombination regions (Figure S3), consistent with the interpretation that gBGC is in part driving the observed pattern.

### Simulations with universal gBGC and an excess of GC→AT mutations recapitulate observed patterns

To evaluate whether the interaction of demographic history with gBGC and a universal imbalance between new GC→AT and AT→GC mutations is sufficient to produce the observed GC% patterns, we performed forward-in-time simulations using SLiM (Haller & Messer, 2023), roughly following an estimated demographic model of human populations (Gravel et al., 2011).

The demographic model included three populations representing African, European, and East Asian populations, with a severe out-of-Africa bottleneck in the non-African lineage followed by rapid population expansion.

To recapitulate the mutation patterns of the human genome, we implemented a universal and constant mutation bias favoring GC→AT mutations (⍰_GC_→AT/⍰_AT_→GC=2.15), based on estimates from human *de novo* mutation data (Kong et al., 2012). At the empirical human genomic GC content of ∼41%, this mutation rate bias results in an excess of GC→AT over AT→GC mutations introduced each generation. In the absence of other forces, this imbalance would drive the genome toward a lower mutational equilibrium GC content of ∼32%. We therefore specified the strength of gBGC such that the expected equilibrium GC content matched the empirical human genome GC% in the ancestral population (Methods). Simulations were initiated at this expected equilibrium genomic GC content and ran for 10Na burn-in generations prior to population split to allow the ancestral population to approach equilibrium in both SFS and stable GC% at segregating sites.

Our simulation results qualitatively mirror our empirical findings in the 1000 Genomes Project data in several aspects. In general, rare variants exhibited higher GC content than common variants (59.4% vs. 53.9%). Specifically, relative to the bottlenecked non-African populations (EUR and EAS), the African population showed lower average DAF and higher GC% at common SNPs (MAF≥5%) (54.83% for AFR vs. 53.58% and 53.01% for EUR and EAS) but higher average DAF and lower GC% at rare SNPs (MAF<5%) (59.33% for AFR vs. 59.44% and 59.45% for EUR and EAS) (Table 2, Figure 4A). These opposing patterns substantially attenuated the inter-population differences when all variants were considered (58.02% for AFR vs. 57.73% and 57.69% for EUR and EAS). As in our empirical observations, relaxing the MAF threshold progressively decreased the difference in GC% between populations (Figure 4B). Most importantly, stratifying variants into non-overlapping, fine-resolution MAF bins revealed a continuous downward trend across all populations and a reversal in the direction of inter-population differences: African populations show lower GC% in rare frequency bins up to ∼12% MAF, whereas non-African populations show lower GC% at higher frequencies (Figure 4C), again recapitulating the empirical pattern (Figure 1C).

**Fig. 4.**
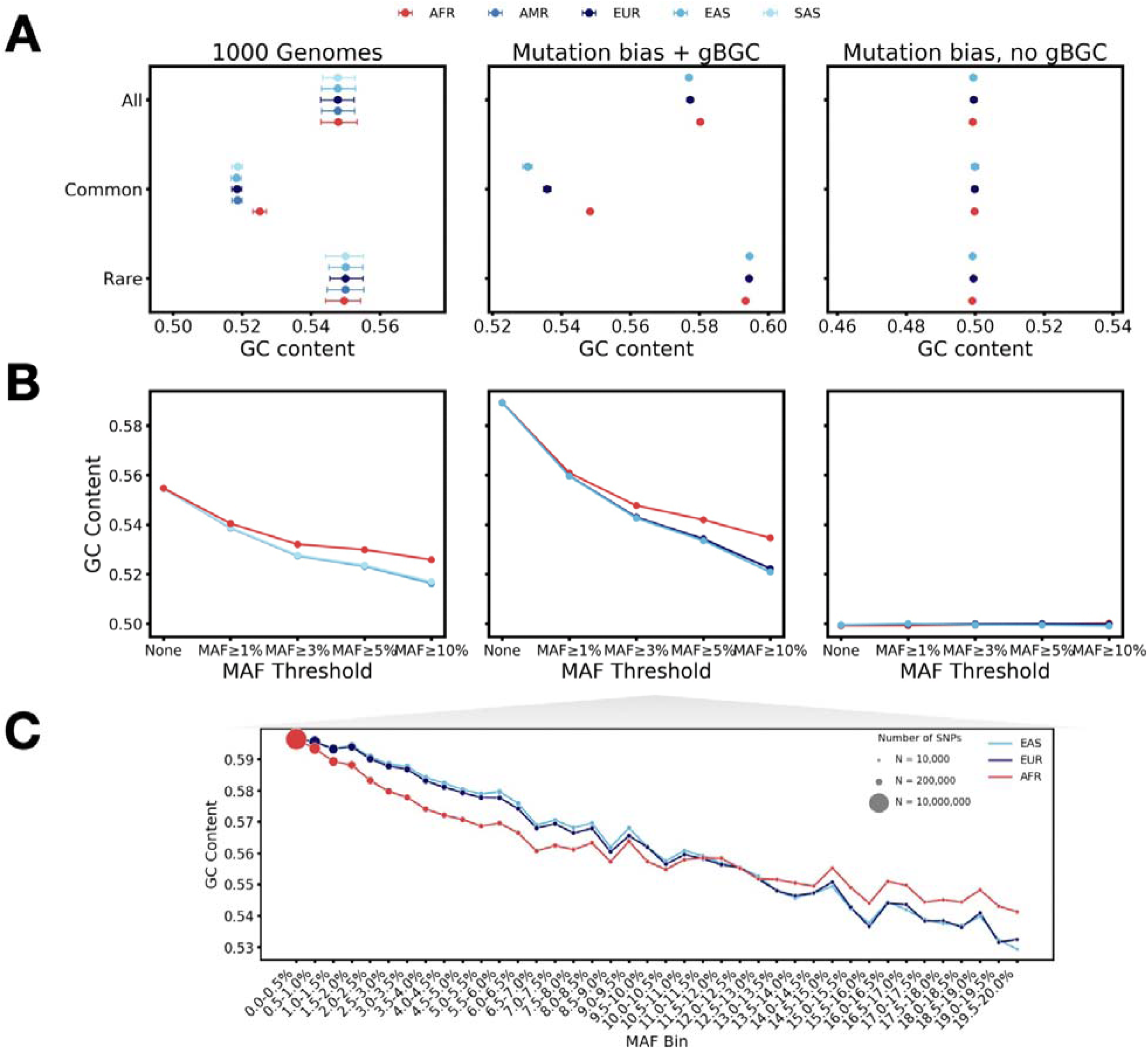
GC content at polymorphic sites from simulating a universal GC→AT mutation bias and GC-biased gene conversion matches observed patterns from 1000 Genomes across the frequency spectrum. **A.** Average GC proportion across polymorphic sites for each population group at common SNPs (MAF≥5%), rare SNPs (MAF<5%), and all SNPs estimated from 1000 Genomes, a simulation scenario with a universal GC→AT mutation bias ⍰GC→AT/⍰AT→GC=2.15 and a gBGC transmission bias = 0.500014 tuned to match the empirical human genome GC% of ∼41%; **B.** GC% across polymorphic sites calculated from different variant sets: No MAF threshold, ≥1%, ≥3%, ≥5%, and ≥10% MAF for each population group in 1000 Genomes and the two simulated scenarios; **C.** Population-specific GC% across SNPs in non-overlapping MAF bins ranging from 0 to 20% (0.5% width) for the mutation bias + gBGC simulation scenario. Marker size is proportional to the number of SNPs in each bin. **Alt text:** Plots comparing GC content at polymorphic sites between populations in different simulation scenarios and in an empirical dataset.

**Table 2.** Average DAF for each population group in our simulation (with ⍰_GC_→AT/⍰_AT_→GC =2.15 and gBGC transmission bias = 0.500014) at GC→AT and AT→GC mutation types along with total AT and GC% across SNPs in different variant bins stratified by MAF: all, common (5%≤MAF≤95%), and rare (MAF<5%).

| All SNPs, n=39,060,136 |  |  |  |  |  |
| --- | --- | --- | --- | --- | --- |
| Mutation Type |  | GC→AT | AT→GC | AT% | GC% |
| SNP Abundance |  | 59.70% | 40.30% |  |  |
| Average DAF |  |  |  |  |  |
| Population | AFR | 8.80% | 8.88% | 41.98% | 58.02% |
|  | EUR | 10.37% | 10.46% | 42.27% | 57.73% |
|  | EAS | 10.59% | 10.68% | 42.31% | 57.69% |
| Common SNPs (MAF≥5%), n=11,201,378 |  |  |  |  |  |
| Mutation Type |  | GC→AT | AT→GC | AT% | GC% |
| SNP Abundance |  | 59.59% | 40.41% |  |  |
| Average DAF |  |  |  |  |  |
| Population | AFR | 25.05% | 25.18% | 45.17% | 54.83% |
|  | EUR | 31.48% | 31.56% | 46.42% | 53.58% |
|  | EAS | 34.47% | 34.55% | 46.99% | 53.01% |
| Rare SNPs (MAF<5%), n=27,858,758 |  |  |  |  |  |
| Mutation Type |  | GC→AT | AT→GC | AT% | GC% |
| SNP Abundance |  | 59.75% | 40.25% |  |  |
| Average DAF |  |  |  |  |  |
| Population | AFR | 2.11% | 2.14% | 40.67% | 59.33% |
|  | EUR | 1.73% | 1.76% | 40.56% | 59.44% |
|  | EAS | 1.63% | 1.67% | 40.55% | 59.45% |

In contrast, in control simulations incorporating the same mutation bias (⍰_GC_→AT/⍰_AT_→GC =2.15) but no gBGC, the genomic GC content is maintained at the expected mutational equilibrium of ∼32%, at which the GC→AT and AT→GC mutation fluxes are balanced. Furthermore, the numbers of GC→AT and AT→GC mutations remain equal across frequency bins within each population, regardless of demographic history. Consequently, GC content at segregating sites remains approximately constant at 50% across allele frequency bins and populations (Figure 4), suggesting that demographic history interacting with mutation bias alone is insufficient to generate the observed empirical patterns. Together, these results demonstrate that under long-term genomic GC content equilibrium, an excess of GC→AT over AT→GC mutations is required to counterbalance gBGC; this mutation imbalance, combined with demographic distortion of the SFS, is sufficient to generate the frequency-dependent GC content patterns observed across populations, without invoking population-specific shifts in the mutation spectrum.

## Discussion

### Reinterpreting GC content variation at polymorphic sites

Previous studies reported lower GC content at common SNPs in bottlenecked or domesticated populations and interpreted this pattern as evidence for rapid shifts in base composition or mutation bias. Our analyses show that this signal is highly dependent on allele frequency conditioning. Although we replicated the previously reported differences at common variants, the direction reversed among rare variants and nearly disappeared when all variants were considered. This observation aligns with prior analyses of the Simons Genome Diversity Project (SGDP) dataset, which found broadly similar accumulation rates of GC→AT and AT→GC mutations across human populations, with the possible exception of the deeply diverged San population not included in the 1000 Genomes dataset analyzed here (Do et al., 2015). Similar attenuation was observed in mouse, maize, and silkworm, indicating that this effect is not unique to recent human demographic history.

These observations can be explained by a simple population genetic mechanism. Bottlenecked populations have distorted site frequency spectra, including higher average derived allele frequencies (DAF) among common SNPs. Because GC→AT mutations outnumber AT→GC mutations among segregating variants, this shift lowers GC content at common SNPs in bottlenecked populations. To illustrate this intuition, consider a set of SNPs in which a fraction are GC→AT mutations and 1 − *x* are AT→GC mutations, with both classes having average DAF of *f*. The average GC content of this SNP set is then *GC*% = *x*(1 − *f*)+ (1 − *x*)*f*= *x* + (1 − 2*x*)*f*. Because GC→AT mutations generally outnumber AT→GC mutations among segregating variants (*x > 0.5*), the GC content is expected to be lower in the bottlenecked population with higher *f.* Moreover, since the fixation probabilities of GC→AT and AT→GC mutations may differ between populations due to demographic influences, inter-population differences in the GC% at polymorphic sites, even across all SNPs (Fig. 2), do not necessarily imply divergence in future GC content.

This same reasoning also explains why rare variants have higher GC content than common variants: rare variants are more dominated by low-frequency GC→AT mutations (i.e., tiny) whose ancestral alleles are GC bases. Importantly, an excess of GC→AT mutations among segregating variants does not imply ongoing genome-wide decline in GC content, as this mutational imbalance can be counteracted by gBGC among mutations that eventually fix, leading to equal amounts of GC→AT and AT→GC substitutions.

Compared to the “mutation explanation” proposed by Li et al (2015), our “demography explanation” is more parsimonious, because it does not invoke inter-population differences in mutation spectrum or gBGC strength. Instead, we assume long-term equilibrium in genomic GC content and the same gBGC transmission bias across populations; under equilibrium, this GC-favoring fixation bias requires an opposing excess of GC→AT over AT→GC mutations, features observed in most species studied to date (Keightley et al., 2009; Hershberg & Petrov, 2010; Lynch, 2010; Näsvall et al., 2023; Bergeron et al., 2023). Forward simulations combining these shared evolutionary processes with population-specific demographic history recapitulated the observed patterns, providing support for sufficiency of this “demography explanation”.

Residual GC content differences across all SNPs and variation among species While relaxing allele frequency thresholds reduced inter-population GC% differences across all species, the effect was most pronounced in humans, where differences reached negligible levels. In contrast, maize and mouse exhibited greater residual signals even when all variants were considered. This cross-species variation in residual inter-population GC% differences may reflect a combination of biological and technical factors, including demographic history (both divergence time and effective population size), recombination landscapes, and sample size, although population-specific evolution in mutation patterns cannot be excluded.

Biologically, these differences could influence the population-scaled strength of gBGC, which determines the degree to which the site frequency spectra of GC→AT and AT→GC variants deviate from neutral expectations. On the other hand, technical constraints such as small sample sizes lack the power to capture the most recent, rare mutations in the population; consequently, the demographic signal at common variants may remain disproportionately represented in the aggregate GC% of all variants, creating apparent differences even in the absence of true evolution in mutation patterns.

The evolutionary scale of the population split may also explain the persistence of the residual signal in some species. Unlike the relatively recent human population divergence, the mouse subspecies diverged approximately 350,000 to 500,000 years ago, representing a significantly deeper divergence in terms of drift units (*T/2Ne*) (Geraldes et al., 2011; Fujiwara et al., 2022). In such diverged lineages, a larger fraction of all variants consists of population private mutations that have evolved under independent mutational, demographic and gBGC regimes for longer times. Consequently, the “all SNP” aggregate in the mouse subspecies could reflect genuine biological divergence in base composition accumulated over time, rather than a transient frequency-shifting artifact of a recent bottleneck. However, such a hypothesis requires more rigorous testing and systematic exclusion of alternative biological (e.g., differences in recombination rates) or technical explanations (e.g., differences in genome assembly quality).

### Effects of allele frequency filters on GC content at polymorphic sites

The practical implication of our analysis is the substantial influence of the MAF threshold on the quantification of GC content at segregating sites (Figure 1). Rare or ultra-rare variants are commonly excluded from genomic analyses to reduce technical artifacts, especially in low-coverage datasets, but our results show that such filtering can strongly bias summaries of base composition at segregating sites. Because frequency-filtered variants are a non-random subset of polymorphisms, their aggregate GC content need not reflect either all segregating variation or genome-wide base composition. As MAF thresholds become more stringent, average GC content declines and subtle inter-population differences can be amplified (Figure 1B).

The flip side of this issue is also important: ultra-rare variants, including singletons, are not themselves representative of either all genomic sites or newly arising mutations, because they are conditioned on recent mutation and on observation as polymorphisms in the sample. The first condition enriches segregating sites for more mutable sites and therefore for GC bases, because GC sites typically have a higher mutation rate than AT sites on average (⍰_G/C_→N>⍰_A/T_→N). The second condition means that variants in the same frequency class, such as singletons or doubletons, can differ in allele age across populations due to differences in demographic history and sample size, and therefore are not directly comparable (see (Ghafoor et al., 2023; Mathieson & Reich, 2017) for discussion of effects of these factors for other reasons). In general, subtle inter-population differences among rare variants should not be interpreted as evidence for shifts in mutation patterns, unless alternative explanations— including differential gBGC, recurrent mutation, and demographic effects—have been adequately considered and ruled out.

## Materials and Methods

### Data sources and filtering

#### Polymorphism

We downloaded unphased autosomal polymorphism data from published sources for four species: human (Byrska-Bishop et al., 2022), mouse (Harr et al., 2016), silkworm (Tong et al., 2022), and maize (Bukowski et al., 2018). We selected these datasets because (1) they were based on whole-genome sequencing and provided unbiased surveys of polymorphisms; (2) the evolutionary relationships of the studied populations were known, allowing for distinction between bottlenecked and non-bottlenecked groups; and (3) adequate numbers of SNPs were detected. For all datasets, we kept only biallelic SNPs. To minimize potential genotyping errors, only SNPs genotyped in more than 80% of samples were considered.

Following Li et al., we operationally defined “common variants” as SNPs with minor allele frequency (MAF) greater than or equal to 5% across samples across all populations, “rare variants” as SNPs with MAF less than 5%, and “all variants” as all SNPs with no MAF threshold. To distinguish between bottlenecked and non-bottlenecked groups, we excluded individuals from clearly admixed populations (i.e., with population labels ‘PUR’,’CLM’,’ACB’,’PEL’,’ASW’) in the 1000 Genomes Project in this analysis. We also identified two individuals from inbred maize populations (‘ZEAxppRDLDIAAPEI-2’ and ‘ZEAhwcRAXDIAAPE’) that clustered with wild individuals based on their proportions of A and C bases at polymorphic sites. We hypothesized that these individuals were admixed and excluded them from subsequent analysis.

### Definition of putatively neutral region

To mitigate the effects of purifying selection, we defined “putatively neutral regions” in our datasets by excluding coding and conserved regions of the genome when available (see “Resources” section below for sources). Specifically, we excluded only coding and phylogenetically conserved regions of the human and mouse genomes. For the maize genome, we only excluded coding regions as annotation of conserved regions was not readily available. We note that our “putatively neutral regions” only mitigate potential effects of direct purifying selection but cannot preclude the effects of background selection, which reduces diversity at neutral sites due to selection at linked regions. We did not apply these filters to the silkworm genome because annotations of coding or conserved regions were not available to our knowledge. For analysis of human polymorphism data, we further intersected the “putatively neutral regions” with the Strict Mask of 1000 Genomes dataset and focused on SNPs in the intersected regions.

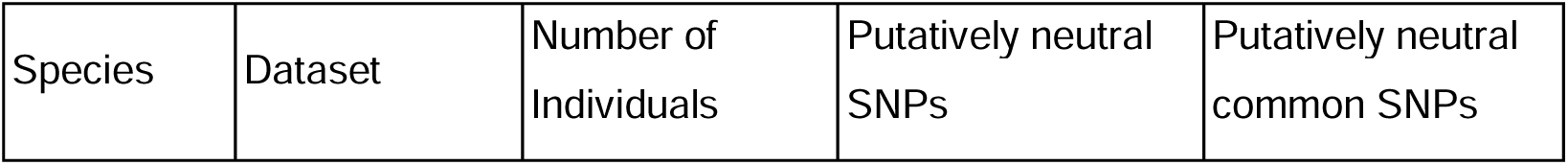
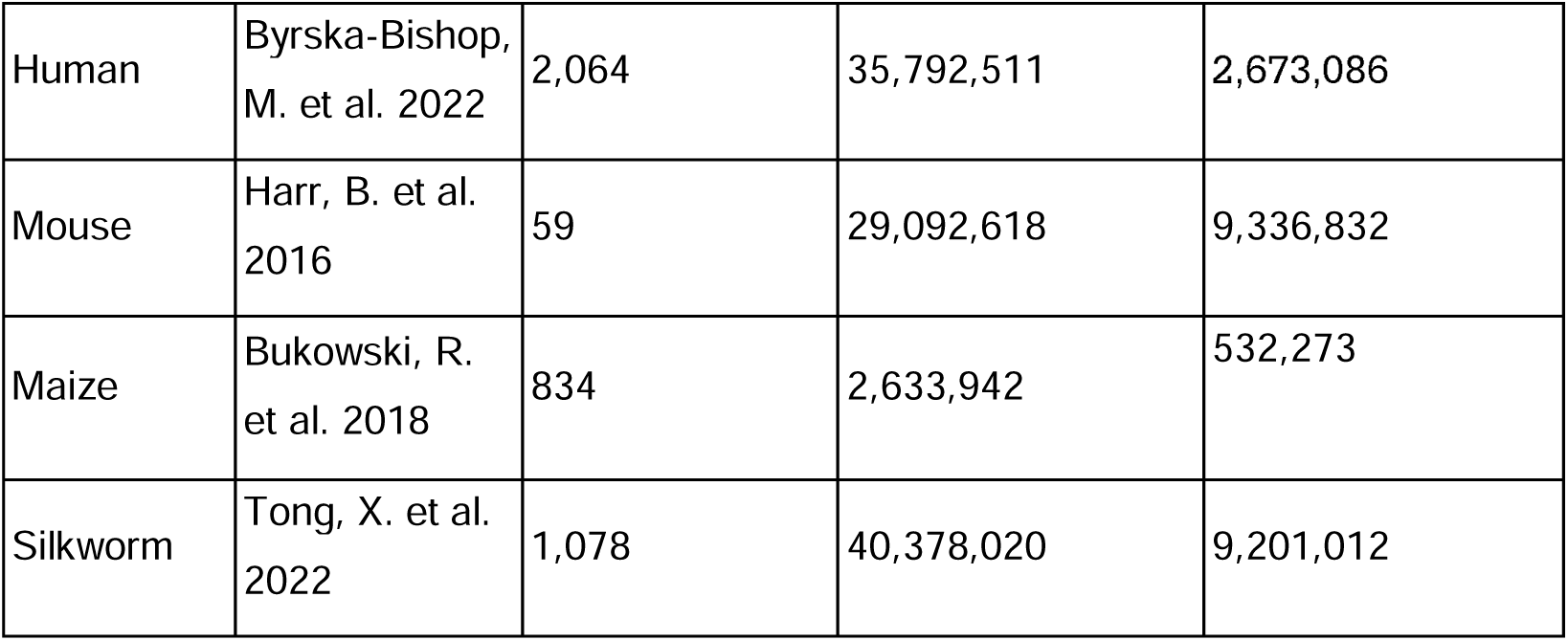

### Resources

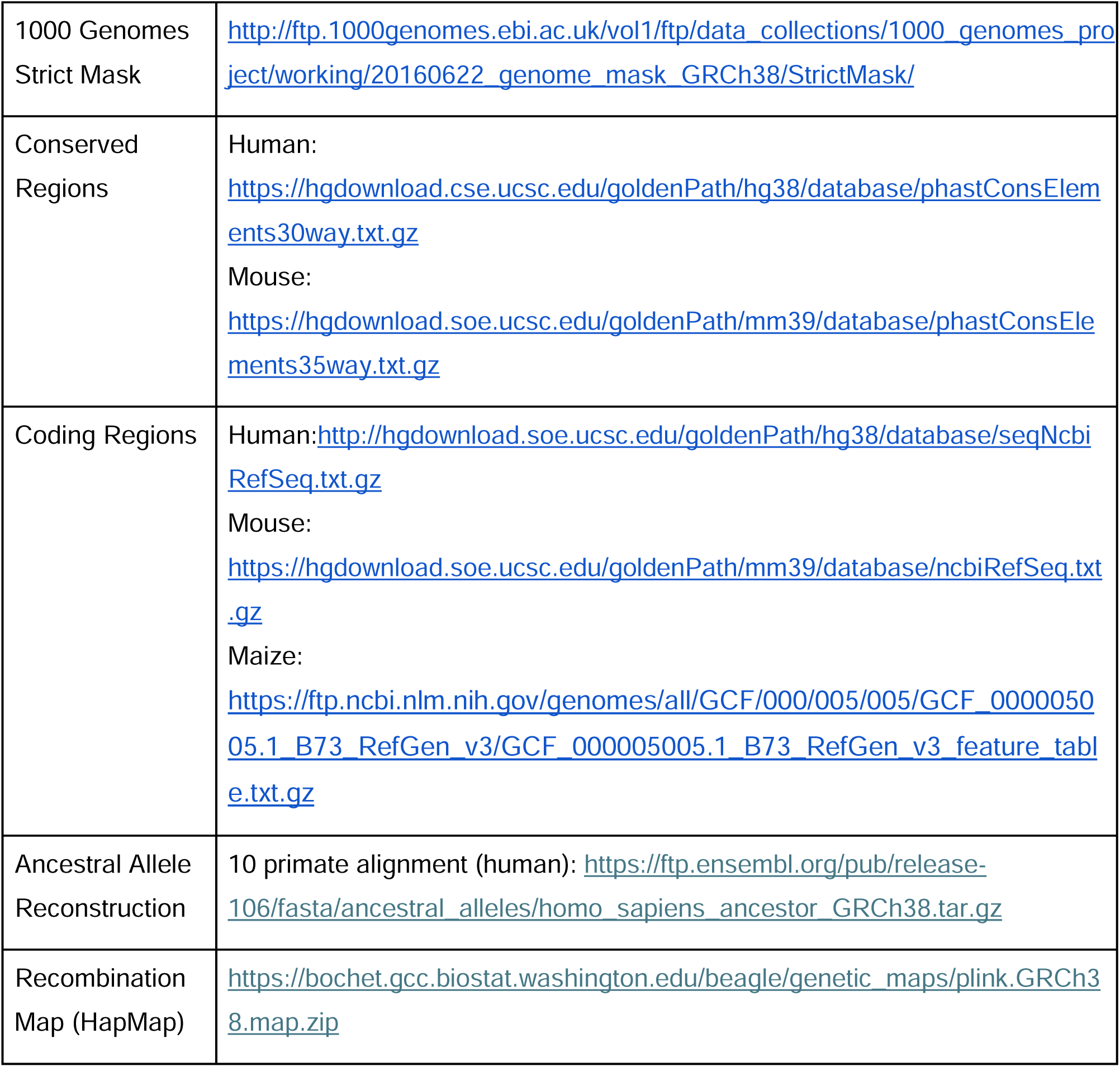

### Base composition across polymorphic sites

Based on the unphased genotype values in the 1000 Genomes Project VCF (0/0 for homozygotes for the reference allele, 0/1 or 1/0 for heterozygotes, and 1/1 for homozygotes for the alternative allele), we counted the number of alternative alleles at each site for each individual. We defined common SNPs (MAF≥5%) and rare SNPs (MAF<5%) based on the global frequency across all 2064 individuals included in this analysis. We then summed the total numbers of A, C, G, and T bases across common SNPs (MAF≥5%), rare SNPs (MAF<5%), and all SNPs for each individual and computed the proportions of A and C alleles on the reference strand (Figure 1A). We used the “Superpopulation” labels assigned to each individual sample in the 1000 Genomes Project to distinguish five continental ancestry populations: African (‘AFR’), European (‘EUR’), American (‘AMR’), East Asian (‘EAS’), and South Asian (‘SAS’). To assess whether GC content at polymorphic sites differed significantly between bottlenecked and non-bottlenecked individuals, we computed per-individual GC content at polymorphic sites as (G+C)/(A+T+G+C) for each SNP set and applied a Wilcoxon rank-sum test to assess differences between populations.

To calculate the average GC% for each population in each SNP set (rare, common, or all), we used population-specific allele frequency values estimated from individual genotype information. Further, we used a block bootstrap approach to estimate uncertainty in GC% across polymorphic sites. Specifically, we partitioned each SNP set into 100 non-overlapping blocks of roughly equal size, randomly sampled 100 blocks with replacement for each bootstrap replicate, and calculated the average GC% across SNPs for each replicate. We performed 500 bootstrap replicates to get the distribution of GC% for each SNP set and estimated 95% confidence intervals from the 2.5th and 97.5th percentiles of the distribution (Figure 4). To further evaluate the effect of allele frequency thresholding on SNP GC%, we applied several global MAF filters (≥1%, ≥3%, ≥5%, and ≥10%) and calculated the GC% for each of the resulting datasets (Figure 1B). Similarly, for each population, we stratified SNPs with 0<MAF<20% into non-overlapping 0.5%-width bins based on the global MAF and computed the average GC% for SNPs in each MAF bin (Figure 1C).

### Ancestral allele polarization

To determine the mutation type that gave rise to each SNP, we used ancestral allele reconstruction based on the 10-primate EPO alignment for humans. For mouse and maize, we assigned ancestral states probabilistically using *est-sfs*, which takes SNPs from a focal species (*Mus musculus; Zea mays*) and an outgroup aligned to the same reference genome (*Mus spretus*; *Tripsacum dactyloides*) to estimate the probability that the major allele of the focal species is the ancestral state (Keightley & Jackson, 2018). For each SNP, we assigned the ancestral allele by a Bernoulli draw weighted by this probability. Based on the inferred ancestral and derived alleles, we then annotated each SNP with the mutational event that gave rise to the polymorphism in the history of the sampled genomes: C>T, C>A, C>G, T>C, T>A, and T>G (where each type includes the corresponding reverse complement; e.g., C>T includes both C>T and G>A).

### Site frequency spectrum analysis

For human, mouse, and maize polymorphisms, we determined the derived allele frequency (DAF) for each SNP using the population-specific allele frequencies computed from individual genotype data (see Data sources and filtering) and the inferred ancestral allele (see “Ancestral allele polarization”). To further investigate the SFS of each population, we calculated the average DAF across all sites (or separately across common and rare variants) of each mutation type. Using the population-specific average DAF values, we determined the average AT and GC content in each mutation type based on the identities of the derived and ancestral alleles. For example, if the average DAF for all SNPs generated by C>T mutations is *X*, the average AT content per individual across these sites would be and GC content (1 − *X*). We used the same block bootstrap approach as described above to generate a distribution of DAF values for each variant set for every mutation type and estimated the 95% confidence intervals as the 2.5th and 97.5th percentiles of the bootstrap distributions (Figure 3B).

### Excluding CpG and TpG sites

Since CpG sites have more than 10x higher transition mutation rates than the genome average, CpG and TpG sites in present-day populations are more likely to have experienced recurrent mutations, which can affect both polarization of the ancestral alleles and quantification of the SFS. For example, two variants at frequencies 1% and 2% due to independent, recurrent mutations at the same site would be treated as one variant at 3%. To evaluate these effects, we computed the SFS for C/T and G/A polymorphisms excluding CpG/TpG/CpA sites. We extracted the 5’ and 3’ flanking nucleotides for each SNP from the GRCh38 reference genome and excluded all sites at which either the reference or the alternative allele, combined with its 5’ or 3’ neighboring nucleotide could form a CpG site. From this filtered dataset, we obtained the SFS by population for different mutation types (Figure S2), which were qualitatively similar to the corresponding results when CpG/TpG/CpA sites were included (Figure 3). We also generated a table of the average DAF for each mutation type for every population group along with the average GC% across SNPs in that variant set and found that patterns of inter-population differences in GC% and average DAF were consistent whether CpG/TpG/CpA sites were excluded or not (Tables S1,S2).

### Regions of low and high recombination rates

Among common SNPs (MAF≥5%), we annotated each SNP with its estimated recombination rate from the HapMap Phase II genetic map (The International HapMap Consortium, 2007), which reflects the long-term sex-averaged recombination landscape of humans. We used the GRCh38 version of this map (see Resources). We looked up the recombination rate for each SNP by interpolation and sorted them to obtain the upper and lower quartiles of the recombination rate distribution (1.33423 cM/Mb and 0.17824 cM/Mb, respectively). This allowed us to categorize the SNPs into two groups: high (roughly 25% of common SNPs with rate ≥1.33423 cM/Mb), and low (∼25% of common SNPs with rate ≤0.17824 cM/Mb) recombination rate regions. We then calculated the reference strand base composition as described earlier (Figure S3).

### Forward-in-time simulations of mutation imbalance and gBGC

To evaluate whether gBGC and the resulting imbalance in the numbers of newly arising AT→GC and GC→AT mutations, coupled with differences in demographic history, are sufficient to reproduce the observed inter-population differences in GC% at polymorphic sites, we performed forward-in-time simulations using SLiM 4.0.1 (Haller & Messer, 2023). We considered two scenarios. In the primary scenario, gBGC was modeled together with the observed AT mutation bias in per-site mutation rate (i.e., ⍰_GC_→AT > ⍰_AT_→GC), maintaining genomic GC content above the mutation-bias-only equilibrium and thereby producing an excess of GC→AT over AT→GC mutations. In the second scenario, AT mutation bias was modeled without gBGC, and genomic GC content was initialized at the corresponding mutation-bias-only equilibrium, at which the opposing mutation fluxes are balanced. We then applied analyses analogous to those used for the empirical data to the simulated populations.

The following demographic and genomic parameters were shared between the two scenarios unless otherwise specified. Each simulation consisted of a 1Mb segment with a per-base per-generation mutation rate of µ=6.89×10⁻ and a recombination rate of 1×10⁻ per base per generation. All mutations were treated as selectively neutral. A total of 2,000 independent replicate simulations were performed under each scenario.

The demographic model followed Gravel et al. (2011) and included three populations: ancestral African (AFR), European (EUR), and East Asian (EAS). The simulation began with an ancestral African population of N = 7,310. After a burn-in period of 10Na = 73,100 generations, the ancestral population expanded to N = 14,475. At generation 76,968, a Eurasian ancestral population (N = 1,861) split from the African population, with symmetric migration rates of 1.52×10⁻ between the two. At generation 78,084, the European (N = 1,032) and East Asian (N = 554) populations split, with the three-way migration rates among AFR, EUR, and EAS set to 2.54×10⁻, 7.77×10⁻, and 3.12×10⁻ respectively. Following the split, European and East Asian populations underwent exponential growth at rates of 0.378% and 0.478% per generation, respectively, until the final generation (79,024), when the simulation was halted and genomes were sampled. To mimic the number of sampled individuals in 1000 Genomes Project, 504 AFR, 503 EUR, and 504 EAS individuals were sampled at the final generation, and their genomes were output as VCF files.

Mutation bias was parameterized via a nucleotide mutation rate matrix in which all AT→GC mutations were assigned a relative rate of 1.0 and all GC→AT mutations were assigned a relative rate of 2.15, yielding ⍰_GC_→AT/ ⍰_AT_→GC= 2.15 (Kong et al., 2012). Under mutation bias alone, this rate asymmetry is expected to produce an equilibrium GC content of ∼32%, at which the lower abundance of GC relative to AT sites compensates for the higher per-site GC→AT mutation rate, such that the opposing GC→AT and AT→GC mutation fluxes are balanced. In the presence of gBGC, however, GC content is maintained above this mutation-bias-only equilibrium, resulting in an excess of new GC→AT over AT→GC mutations (i.e., mutation imbalance).

GC-biased gene conversion (gBGC) was implemented using SLiM’s gene conversion model with a non-crossover fraction of 0.7, a mean tract length of 1,500bp, and a GC transmission bias of 0.500014. We selected this value of gBGC transmission bias parameter so that the predicted equilibrium GC content under the combined effects of mutation bias and gBGC matched the empirical human genome-wide GC% of ∼41% (International Human Genome Sequencing Consortium et al., 2001), which is higher than the ∼32% equilibrium expected under mutation bias alone. In other words, by maintaining GC content above the mutation-bias-only equilibrium, gBGC sustains the mutation imbalance described above.

We derived the gBGC strength parameter, B, required to match the empirical human genome GC content as follows. At equilibrium, the ratio of GC to AT content equals the ratio of the AT→GC to GC→AT fixation probabilities weighted by their respective mutation rates:

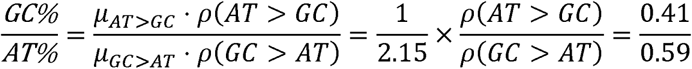

which requires a fixation probability ratio of:

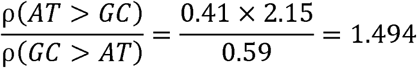

Under gBGC, with a gBGC selection coefficient of 2*b*, where *b* is the conversion bias, the fixation probabilities of AT→GC and GC→AT mutations are given by:

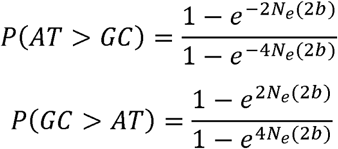

If we define the population-level gBGC strength as 4 = 4*Neb*, the fixation probability ratio can be reduced to:

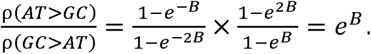

Setting *e^B^* = 1.494 gives *B* = 0.4015; plugging in the ancestral effective population size

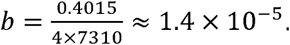

The SLiM gene conversion bias parameter corresponds to 0.5 + *b*, yielding a value of 0.500014, which was used in the mutation-imbalance-plus-gBGC simulation. Our estimate of genome-wide average gBGC strength *B* = 0.4015 aligns well with previous estimates based on derived allele frequency distribution in humans, which generally place global gBGC intensity in the nearly neutral range (e.g. B≈0.38-0.43, (Glémin et al., 2015); B≈0.25-0.75 depending on mutation type and population (Bergman & Heide Schierup, 2021)). We further confirmed that genomic GC% remained unchanged during the simulation by calculating nucleotide frequencies at (1) the start of the burn-in period, (2) the end of the burn-in period, and (3) the end of the simulation. At every stage, GC% was stably maintained at expected equilibrium.

For the control scenario, we simulated mutation bias (i.e., ⍰_GC_→AT > ⍰_AT_→GC) without gBGC. All parameters were identical to those in the “mutation bias + gBGC” scenario described above, with two exceptions: the gene conversion model was omitted entirely, and the ancestral sequence was initialized at the corresponding mutation-bias-only equilibrium GC content of ∼32% (nucleotide frequencies of 0.159 for G and C and 0.341 for A and T). In this scenario, the lower abundance of GC relative to AT sites perfectly compensates for the higher per-site GC→AT mutation rate, such that the numbers of newly arising GC→AT and AT→GC mutations are balanced. Thus, mutation-rate bias alone does not produce an imbalance in mutation counts when the genome is at its corresponding mutation-bias equilibrium.

From each replicate’s VCF output, we repeated the analyses analogous to those performed on the empirical data for Fig. 1 and Table 1. We used global MAF across samples from all three populations to assign SNPs to SNP sets (e.g. rare MAF<0.05; common MAF≥0.05, and all), ensuring consistent SNP set membership across populations (Fig. 4). For calculating average DAF within different SNP sets (Table 2), we used population-specific allele frequencies. Within each SNP set/bin, we computed: (1) mean DAF separately for GC→AT and AT→GC mutations in each population; and (2) average GC% for each population, as described earlier. 95% confidence intervals were estimated using the block bootstrapping procedure as described earlier.

## Acknowledgements and funding sources

We thank Iain Mathieson and other members of the Gao and Mathieson laboratories for helpful discussions. This work is supported by a Research Fellowship (FG-2021-15702) from the Alfred P. Sloan Foundation (https://sloan.org/) and a grant (R35GM146810) from the National Institute of General Medical Sciences to ZG.

## Data Availability Statement

No new empirical data were generated in support of this research. Data used for this project was obtained from publicly available sources, with specific links to download provided in the Materials and Methods section under Resources.

## Notes

### Competing Interest Statement

The authors have declared no competing interest.

### Summary of Updates

This version of the manuscript has been revised to better situate our findings in the context of previous literature, reduce jargon, and more explicitly acknowledge limitations of our current approach. We also include an additional analysis (Figure 1C and 4C) to further establish the novelty of our findings and strengthen our interpretation.

