## Supplementary Figures for "Interpreting GC content differences across populations at polymorphic sites"

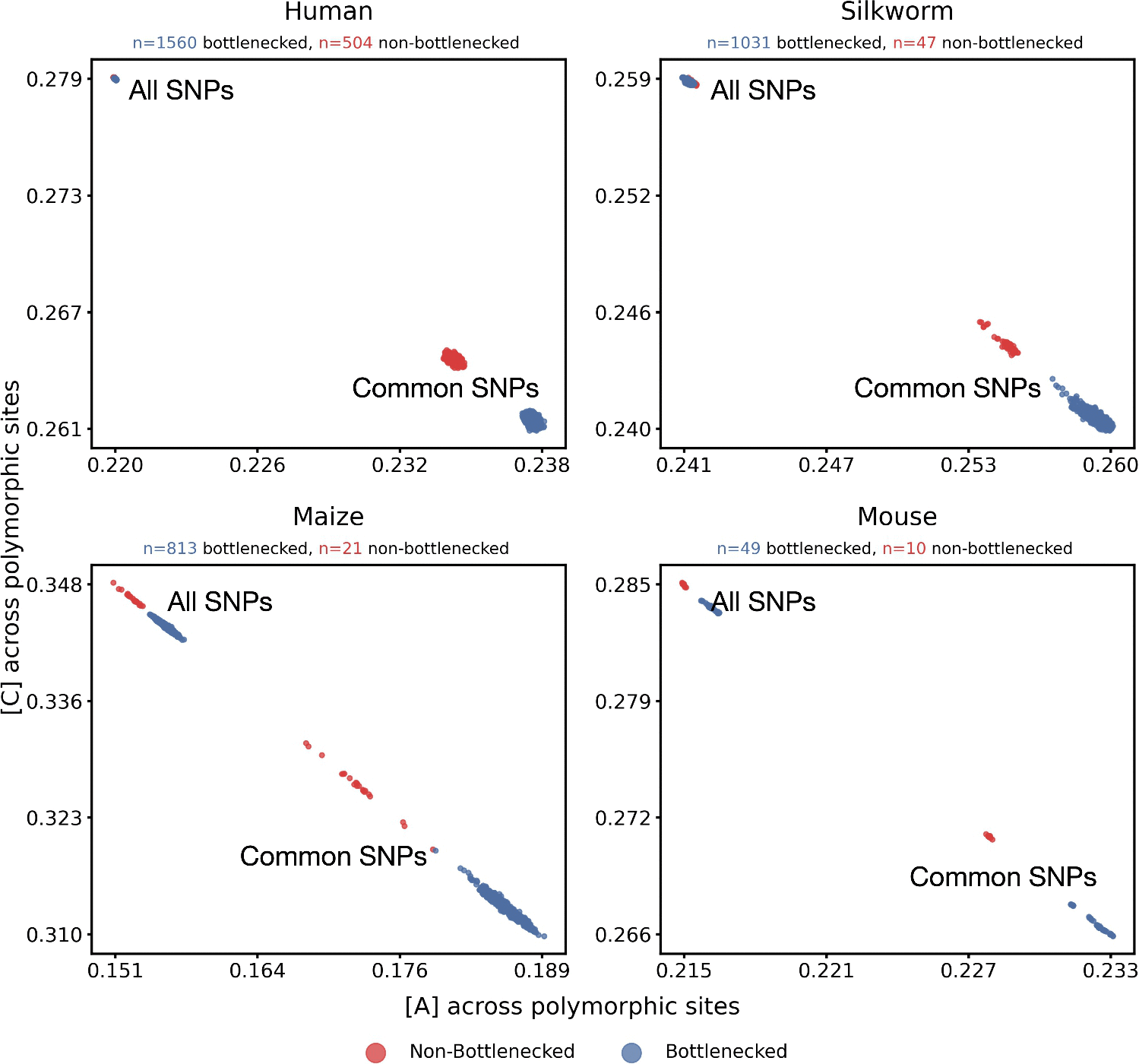

**Supplementary Figure 1. Decrease in inter-population differences in GC% at SNPs is observed even after excluding singletons.** Reference strand base composition plotted as proportion of A vs. proportion of C across common polymorphic sites (MAF≥5%) and all polymorphic sites (no MAF threshold, excluding singletons) between bottlenecked and non-bottlenecked populations in human, mouse, maize, and silkworm.

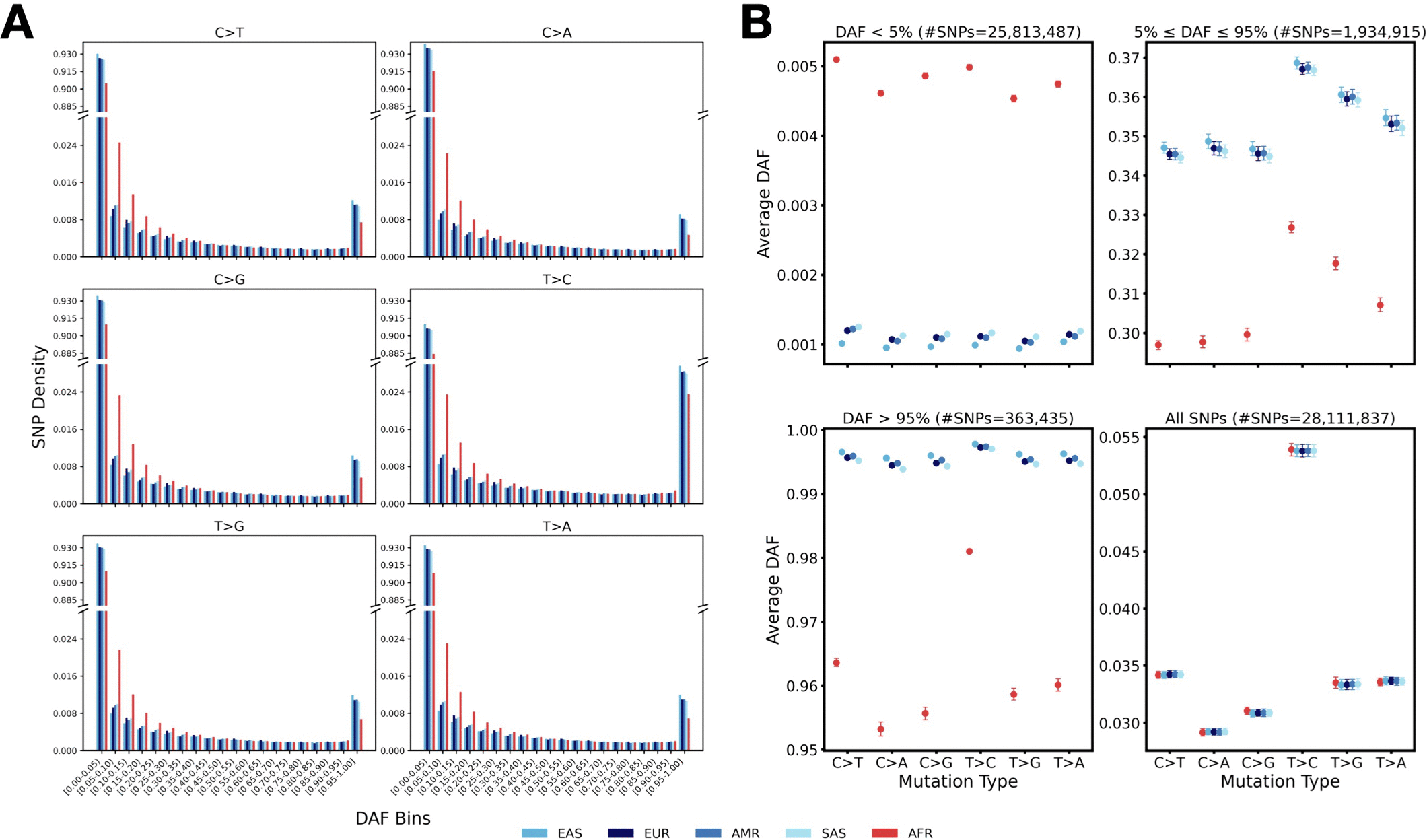

**Supplementary Figure 2. Site frequency spectrum (SFS) and average derived allele frequencies (DAF) for SNPs generated by different mutation types after excluding CpG/TpG/CpA sites. A.** SFS for SNPs of each mutation type across five human population groups. The y-axis is broken between 0.024 and 0.885 to facilitate visualization of differences at both rare and common variants. **B.** Average DAFs for each population group across different variant sets stratified by DAF: DAF<5%, 5%≤DAF≤95%, DAF>95%, and all SNPs, for each mutation type. Error bars represent 95% confidence intervals estimated using block bootstrap (see Methods). Each population group is represented by a unique color, with AFR in red and non-AFR groups in different shades of blue. All CpG/TpG/CpA sites are excluded.

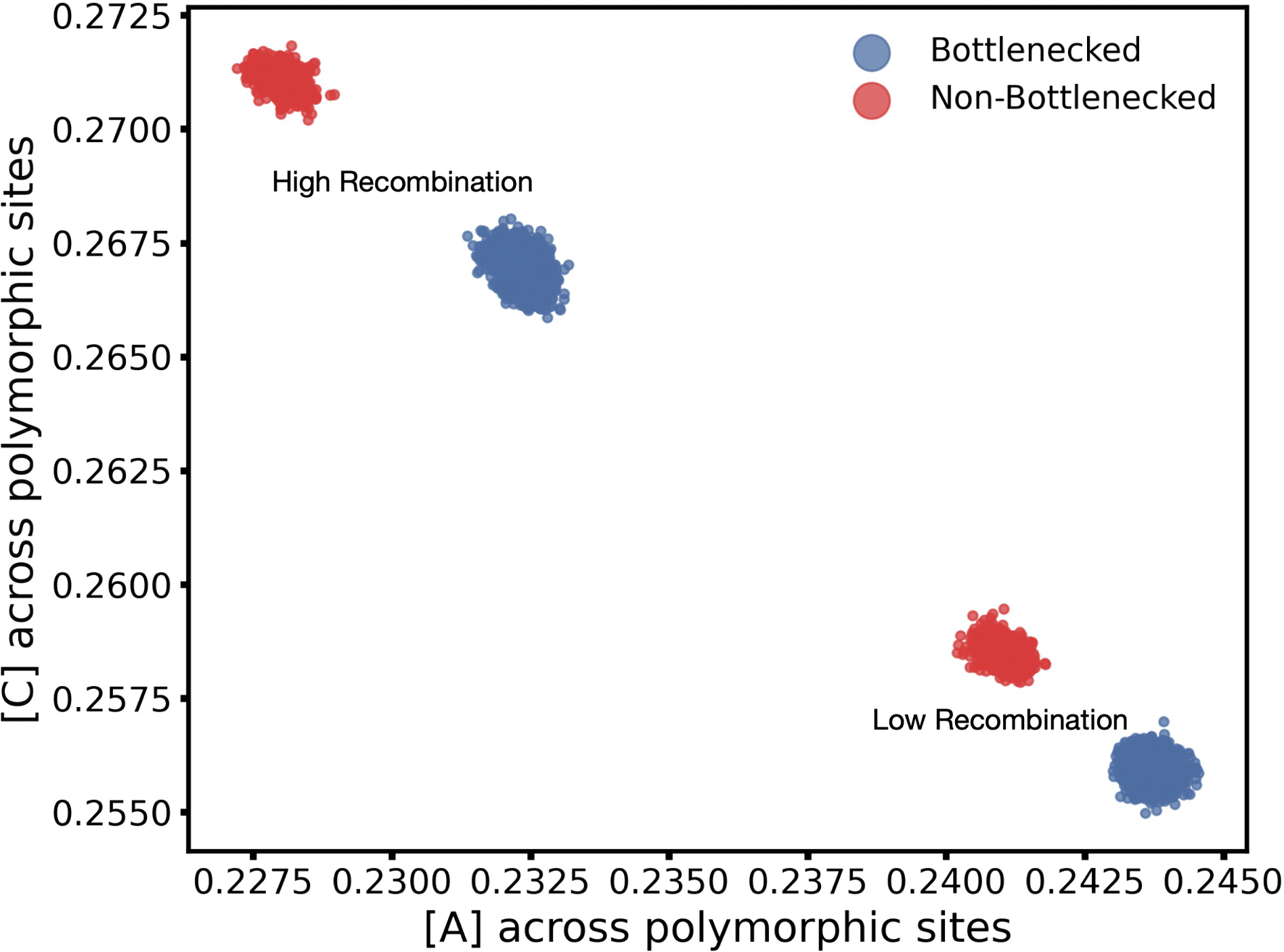

**Supplementary Figure 3. Patterns of inter-population difference in [C] and [A] reflect effects of gBGC on common SNPs.**

Reference strand base composition plotted as proportion of A vs. proportion of C across common polymorphic sites (MAF≥5%) in low recombination rate regions (SNPs with rate ≤0.17824 cM/Mb) and high recombination rate regions (SNPs with rate ≥1.33423 cM/Mb) in bottlenecked (non-AFR) and non-bottlenecked (AFR) population groups.
